# Adaptive divergence of meiotic recombination rate in ecological speciation

**DOI:** 10.1101/2020.07.24.219576

**Authors:** Swatantra Neupane, Sen Xu

**Author notes:** Correspondence author: Sen Xu.

## Abstract

Theories predict that directional selection during adaptation to a novel habitat results in elevated meiotic recombination rate. Yet the lack of population-level recombination rate data leaves this hypothesis untested in natural populations. Here we examine the population-level recombination rate variation in two incipient ecological species, the microcrustacean *Daphnia pulex* (an ephemeral-pond species) and *D. pulicaria* (a permanent-lake species). The divergence of *D. pulicaria* from *D. pulex* involved habitat shifts from pond to lake habitats as well as strong local adaptation due to directional selection. Using a novel single-sperm genotyping approach, we estimated the male-specific recombination rate of two linkage groups in multiple populations of each species in common garden experiments and identified a significantly elevated recombination rate in *D. pulicaria*. Most importantly, population genetic analyses show that the divergence in recombination rate between these two species is most likely due to divergent selection in distinct ecological habitats rather than neutral evolution.

**Significance statement:** Whether directional selection during adaptation to a novel habitat results in elevated meiotic recombination remains largely untested in natural populations. This work examines the population-level recombination rate in two closely related microcructacean species *Daphnia pulex* and *D. pulicaria* using single-sperm genotyping approach. Recombination rate data from two linkage groups show elevated recombination rates in *D. pulicaria* whose divergence from *D. pulex* is accompanied by a habitat shift. Importantly, population genetic analysis suggests that this divergence of recombination is likely adaptive rather than neutral.

## Introduction

Meiotic recombination is a hallmark of meiosis as it occurs in the majority of sexually reproducing eukaryotes (Otto & Lenormand 2002; Cavalier-Smith 2002). Although it remains contested as to why recombination originated in the last common ancestor of eukaryotes (Cavalier-Smith 2002; Kondrashov 1988), recombination plays an essential role in repairing the actively induced double-strand DNA breaks in the prophase I of meiosis (Pâques & Haber 1999). The presence of at least one recombination event (i.e., crossover event) per chromosome arm between homologous chromosomes ensures the correct segregation of chromosomes into daughter cells, preventing chromosome non-disjunction and aneuploidy (Hassold & Hunt 2001).

Besides its well-known role in creating new haplotypes and in facilitating adaptation (Rice 2002), meiotic recombination is an important evolutionary force shaping the eukaryotic genomic architectures. Recombination rate is a determinant of the distribution of genetic diversity in the genomes (Lercher & Hurst 2002; Booker *et al*. 2017; Begun & Aquadro 1992; Charlesworth *et al*. 1993). Recombination reduces selection interference between linked sites (Hill & Robertson 1966; Barton 1995b; Cutter & Payseur 2013; Felsenstein & Yokoyama 1976) and slows down the accumulation of deleterious mutations and of transposable elements (Lynch *et al*. 1993; Dolgin & Charlesworth 2008; Rizzon *et al*. 2002; Kent *et al*. 2017). Moreover, recombination and associated biased gene conversion can influence codon usage bias (Comeron *et al*. 1999; Pouyet *et al*. 2017) and base composition (Duret & Arndt 2008; Mugal *et al*. 2015).

Meiotic recombination rate varies greatly at multiple biological levels, e.g., within genome, between individuals and populations, and between species (Dapper & Payseur 2017; Ortiz-Barrientos *et al*. 2016; Ritz *et al*. 2017; Smukowski & Noor 2011). Understanding the genetic basis and evolutionary forces underlying such variation is a major challenge to biologists. Striking progress has been made in mapping the genetic factors responsible for within-genome variation and for between-individual variation. For example, the zinc finger domain protein PRDM9 is a major determinant of recombination hotspots in the genomes of human and mice (Baudat *et al*. 2010; Myers *et al*. 2010; Brick *et al*. 2012; Grey *et al*. 2011). Also, promoters and transcription start sites have been identified to be associated with elevated recombination rate in dogs (Auton *et al*. 2013), the yeast *Saccharomyces cerevisiae* (Pan *et al*. 2011), birds (Singhal *et al*. 2015), and *Arabidopsis* (Choi *et al*. 2013). On the individual level, several meiosis-related genes (e.g., *Rnf212, Cplx1, Rec8, Prdm9*) have been identified to be responsible for variation of recombination rates in mammalian species including humans, cattle, and Soay sheep (Chowdhury *et al*. 2009; Halldorsson *et al*. 2019; Johnston *et al*. 2016; Kong *et al*. 2008; Sandor *et al*. 2012). However, it should be noted that these loci explain only a small portion (∼3-11%) of the phenotypic variance between individuals (Johnston *et al*. 2016; Kong *et al*. 2014).

In contrast, the genetic factors governing the inter-specific variation of recombination rate remains understudied (Dapper & Payseur 2017), although many studies have compared recombination rate differences between closely related species at different genomic scales (Smukowski and Noor, 2011). We note that this research area has drawn increasing amount of attention, with a dicistronic gene mei-217/mei-218 recently identified to be responsible for recombination rate difference between *Drosophila melanogaster* and *D. mauritiana* (Brand *et al*. 2018) andTEX11 and other genes involved in synaptonemal complex suggested as candidates driving the evolution of recombination rate in mammals (Dapper & Payseur 2019). On the other hand, another equally understudied question is whether natural selection plays a role in shaping the between-species divergence. Despite numerous evolutionary theories have examined how natural selection can modulate the evolution and divergence of recombination rates between species, the lack of in-depth population-level data (see below) leaves these theories untested in natural systems, severely limiting our understanding of the evolutionary forces driving recombination rate divergence.

In populations undergoing divergence and incipient speciation, recombination rates could be driven to increase if the breakdown of overrepresented association of alleles, i.e., linkage disequilibrium, is beneficial. Generally speaking, three different situations can lead to the buildup of linkage disequilibrium and determine how recombination rates respond to natural selection. In the presence of linkage disequilibrium caused by weak negative epistasis selection favors increased recombination in a large population in stable environment (Otto & Lenormand 2002). Genetic drift can also lead to the accumulation of linkage between beneficial alleles and deleterious alleles in finite populations, and the increase of recombination rate would be favored by selection to bring together beneficial alleles (Otto 2009). Furthermore, temporal fluctuations in the environment favor different combinations of alleles, which could lead to increased recombination rate in environments with rapid and consistent temporal variation (Otto & Michalakis 1998; Barton 1995a; Charlesworth 1976), whereas in the absence of fluctuations recombination is selected against.

Despite the diverse views on the relative importance of these evolutionary forces in shaping the evolution of recombination rate, it is consistently predicted that transition to a novel environment would lead to an increase of recombination rate due to directional selection (Butlin 2005). Empirical work on indirect selection of physiology-related traits in *Drosophila* supports this view (Korol & Iliadi 1994; Aggarwal *et al*. 2015). However, for domesticated animals that underwent strong directional selection, there seems to be no increase of recombination rate (Munoz-Fuentes *et al*. 2015), contradicting previous views of elevated recombination in domesticated plants (Ross-Ibarra 2004) and animals (Burt & Bell 1987; Poissant *et al*. 2010).

Notably, few studies have directly addressed whether habitat shift in natural populations results in elevation of recombination rate. A key challenge is that, for model organisms where recombination is heavily investigated, e.g., human, mice, *Drosophila*, and yeast, little is known about the ecological changes involved in speciation. Thus, the inter-specific difference between *Drosophila* species, e.g., ∼2-fold difference between *D. melanogaster* and *D. mauritiana* (Brand *et al*. 2018), and the difference between yeast species, e.g., 40% lower recombination rate in *Saccharomyces paradoxus* than in *S. cerevisiae* (Liu *et al*. 2019), are unfortunately de-coupled from the consideration of ecology.

Another challenge in understanding the relationship between ecological shifts, directional selection, and recombination rate evolution is that multi-population data on recombination rate is largely lacking (but see Saleem *et al*. 2001 and Samuk *et al*. 2020). Recombination rate is laborious to measure, which usually involves producing and genotyping hundreds of recombinant progenies with a large number of genetic markers to generate only a single genetic map. Such practice is difficult to scale up to population-level studies. Thus, current estimates of recombination rates for most species are derived from the average recombination rates in the two lineages used for crossing-based map construction. Often the number of genetic maps for a single species remains below a handful except for some heavily studied model organisms and economically important crops and animals, yielding low statistical power for rigorously investigating the driving forces of inter-specific differentiation of recombination rate in a population-genetic framework.

If a genetic map is constructed using computational methods based on linkage disequilibrium through population sequencing, we can obtain estimates of population recombination rates (4·Ne·c), which is an average of two sexes over large span of evolutionary time (McVean *et al*. 2004). The fact that population recombination rate is scaled by effective population size (Ne) makes it difficult to directly estimate recombination rate and can confound comparisons between diverging populations that may have distinct demographic histories (Rogers 2014; Dapper & Payseur 2018). Promising advances to incorporate demographic changes into this approach have emerged in recent years (Kamm *et al*. 2016; Spence & Song 2019). However, studies using this method to compare multiple populations remain rare beyond the classic models, mostly with a focus on intragenomic variation (e.g., Auton *et al*. 2012). Therefore, we argue that the lack of understanding on the population- and individual-level variation of recombination rates ought to be addressed if we are to dissect the genetic basis of recombination rate variation.

The emergence of novel genomic sequencing techniques such as whole-genome sequencing of single-sperm cells (Xu *et al*. 2015) provides an efficient solution to estimating population-level recombination variation (albeit it only measures male-specific recombination rate). Taking advantage of this approach to investigate how ecological shifts and directional selection impact recombination rate, this study examines the male-specific recombination rate in two ecologically distinct, incipient microcrustacean species *Daphnia pulex* and *D. pulicaria*.

A well-known characteristic of the *Daphnia* system resides in its cyclically parthenogenetic reproduction. Under favorable environmental conditions female *Daphnia* produces directly developing embryos (i.e., live birth of neonates released from brood pouch) via apomictic parthenogenesis, generating genetically identical, diploid daughters. However, stressful conditions, e.g., food shortage (Deng 1996) and decrease in temperature, triggers *Daphnia* females to switch to sexual reproduction and also to parthenogenetically produce males via environmental sex determination (Olmstead & Leblanc 2002). The parthenogenetic production of males allows us to amass a large number of males and sperm cells of the same genotype to examine recombination rate.

As members of the *D. pulex* species complex, *D. pulex* and *D. pulicaria* are estimated to have started diverging from 800,000 – 2,000,000 years ago (Colbourne & Hebert 1996; Cristescu *et al*. 2012; Omilian & Lynch 2009). These two species are morphologically nearly indistinguishable (Brandlova *et al*. 1972) but occupy distinct, overlapping freshwater habitats in North America, with *D. pulex* mostly living in ephemeral fishless ponds and *D. pulicaria* inhabiting stratified permanent lakes. Importantly, population genetic data suggest that the divergence of *D. pulicaria* from *D. pulex* most likely involved a habitat transition event from pond to lake systems (Cristescu *et al*. 2012).

As stratified permanent lakes and ephemeral ponds pose distinct selection regimes (e.g., distinct predators, environmental factors), these two species have most likely undergone strong local adaptation and divergent selection in their distinct habitats, resulting in clear physiological and behavioral differences. For example, compared to *D. pulicaria, D. pulex* grows faster to a larger size, reproduces at an earlier age. Also, *D. pulicaria* exhibits diurnal vertical migration in lakes, whereas *D. pulex* displays no such behavior. Interestingly, the frequency of sexual reproduction is also different between the two. *Daphnia pulex* goes through sexual reproduction producing resting eggs before ponds dry up in early summer every year, whereas *D. pulicaria* can persist in lakes largely without sex for a few years (Cáceres & Tessier 2004; Dudycha 2004; Dudycha & Tessier 1999). Notably, prezygotic isolation has developed between these two species (Deng 1997), with *D. pulex* switching to sexual reproduction at long-day hours (16 hours/day) and *D. pulicaria* switching to sexual at short-day hours (10 hours/day). Despite these differences, *D. pulex* and *D. pulicaria* can still generate fertile cyclically parthenogenetic F_1_ offspring in laboratory crossing experiments, indicating the absence of complete reproductive isolation (Heier & Dudycha 2009).

In this pilot study we examine whether neutral evolution (i.e., genetic drift) is sufficient to explain the divergence of meiotic recombination rate between these two species, with the alternative hypothesis being that directional selection involved in ecological shifts better explains the between-species divergence. As our pilot experiment we estimated recombination rate for a 1.5-Mbp on linkage group 8 and a 0.5-Mbp region on linkage group 9 in three geographically isolated populations of each species. Most interestingly, our results yield strong support for significantly higher recombination rate in *D. pulicaria* than in *D. pulex*, and the between-species divergence in recombination rate cannot be accounted for by genetic drift and is most likely due to directional selection.

## Results

### Recombination rate estimates

We performed microsatellite genotyping on whole-genome amplified single-sperm cells to estimate recombination rates for a 1.5-Mbp on linkage group 8 and a 0.5-Mbp region on linkage group 9 in three populations of *D. pulex* and *D. pulicaria* each. Our recombination rate estimates show that *D. pulicaria* tends to recombine at a higher rate than *D. pulex* (Figs. 1 and 2). For the region on linkage group 8 alone, although the mean recombination rate is higher in *D. pulicaria*, no statistically significant difference (t-test P=0.10) is found between the mean of *D. pulex* (16.9 cM, SD=8.4) and that of *D. pulicaria* (28.4 cM, SD=4.5). However, for the region on linkage group 9, the average recombination rate of *D. pulicaria* (mean=24.5, SD=1.4) is higher (t-test P=0.046) than that of *D. pulex* (mean=13.0, SD=6.8). When we compared the recombination rates of both linkage groups in these two species, *D. pulicaria* has an overall significantly higher recombination rate than *D. pulex* (t-test P=0.006).

**Figure 1.**
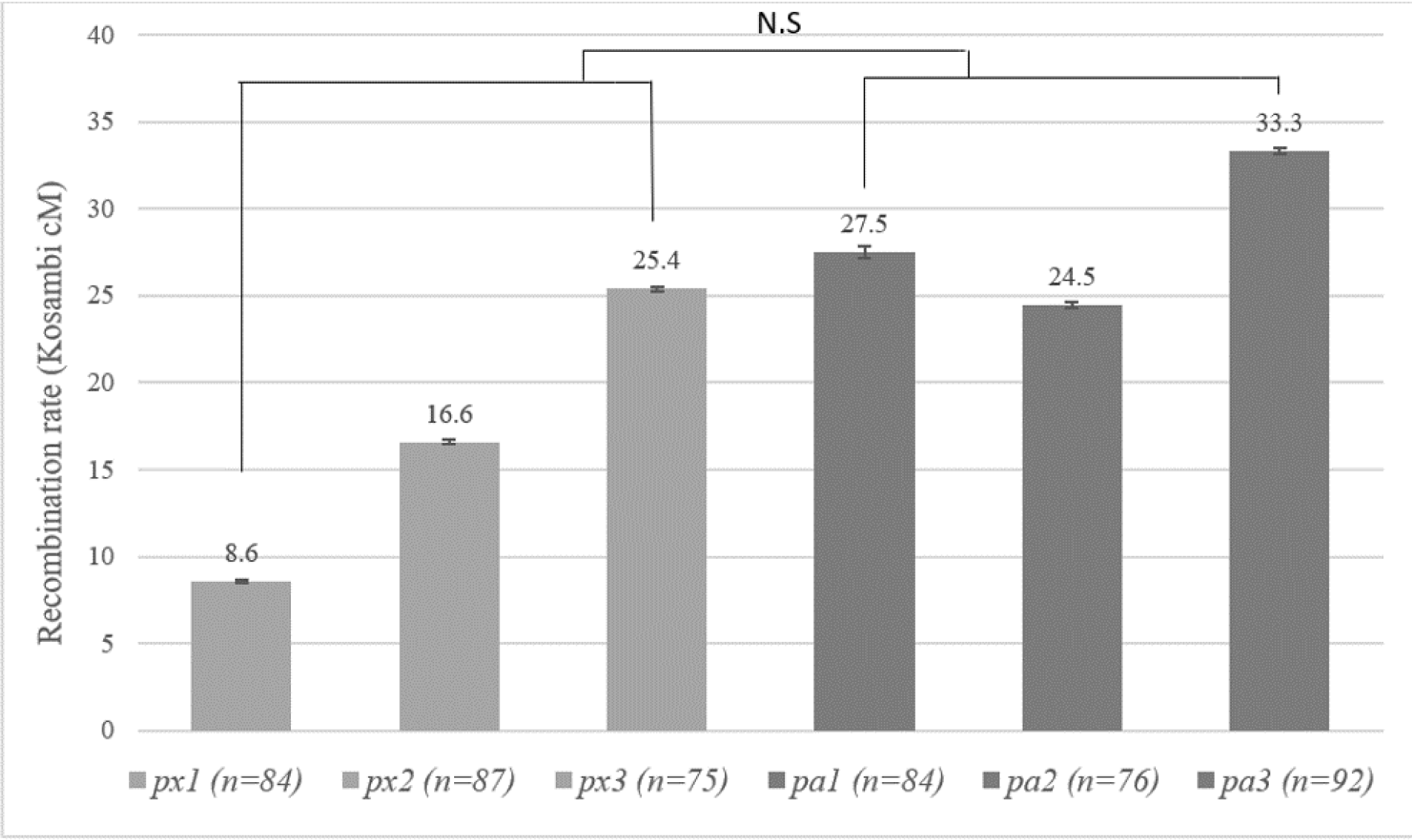
Estimated recombination rates for a 1.5-Mbp region on linkage group 8 for three isolates of *D. pulex* (px) and *D. pulicaria* (pa) each (n represents the number of genotyped sperm). Each grey bar represents the recombination estimate from a specific *Daphnia* isolate with error bar representing standard error. The average recombination rate between these two species is not significantly different (N.S: Not Significant).

**Figure 2.**
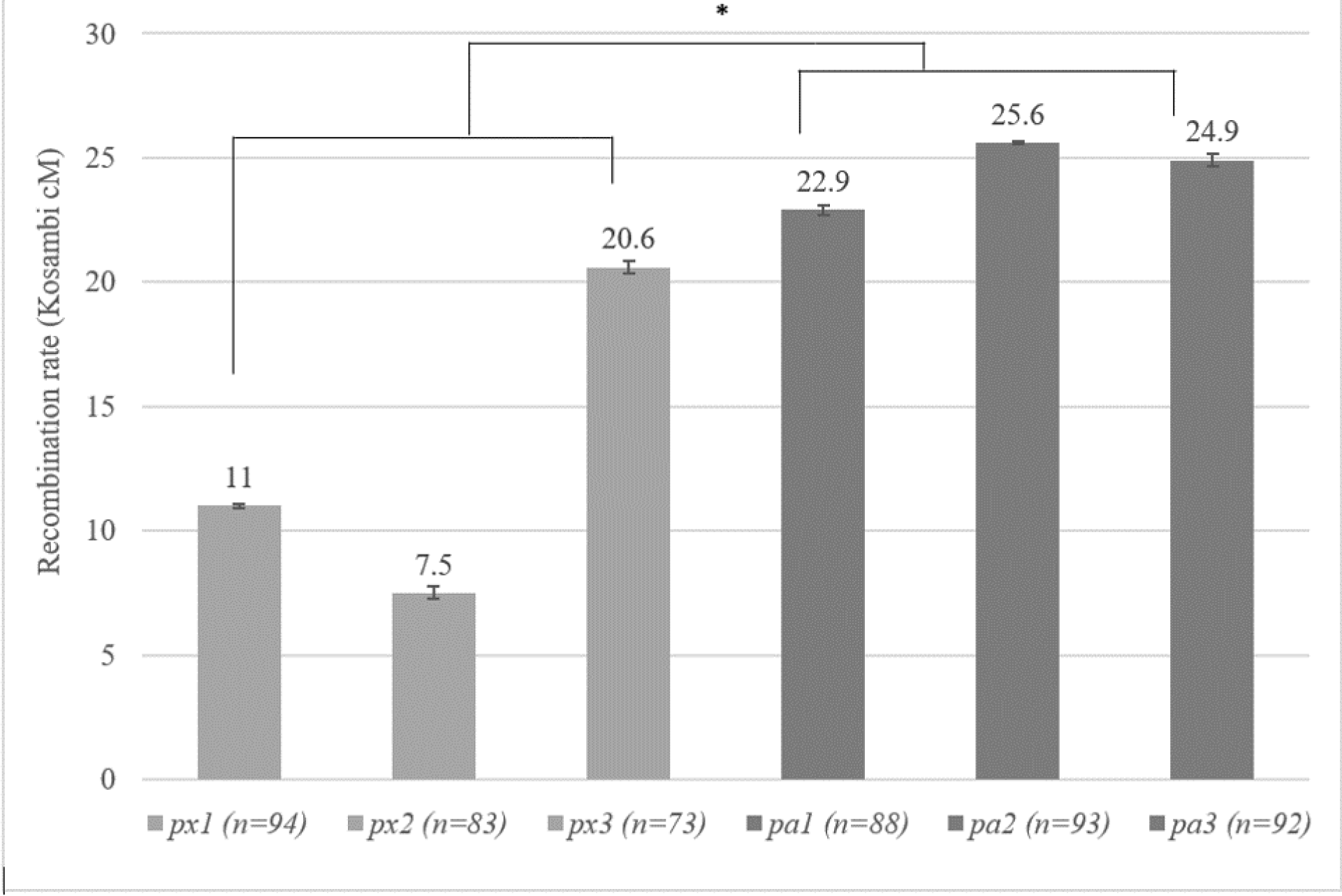
Estimated recombination rates for a 0.5-Mbp region on linkage group 9 for three isolates of *D. pulex* (px) and *D. pulicaria* (pa) each (n represents the number of genotyped sperm). Each grey bar represents the recombination estimate from a specific *Daphnia* isolate with error bar representing standard error. The average recombination rate between these two species is significantly different (* p < 0.05).

Remarkably, our recombination rate estimates show that the within-species recombination rate variation is markedly higher in *D. pulex* than in *D. pulicaria* for regions on both linkage groups. In *D. pulex*, recombination rates of the three sampled populations range from 8.6 Kosambi cM to 25.4 Kosambi cM for the 1.5-Mbp region on linkage group 8 with a nearly 3-fold difference (Fig. 1). On the other hand, within-species variation in *D. pulicaria* for the interval in linkage group 8 is much lower, ranging from 24.5 to 33.3 Kosambi cM among the three examined populations (Fig. 1).

A resembling pattern of distinct within-species variation is also observed for linkage group 9. For *D. pulex*, the map distance of the 0.5-Mbp region on linkage group 9 range between 7.5 cM and 20.6 Kosambi cM, showing a nearly 3-fold difference among populations (Fig. 2). However, for *D. pulicaria*, the recombination rates of the same interval in linkage group 9 from the examined populations show little variation between 22.9 and 25.6 Kosambi cM (Fig. 2).

### P_st_-F_st_ comparison

An important approach for determining whether the divergence of phenotypic traits is neutral is to compare Q_st_ of phenotypic traits and F_st_ of neutral molecular markers. As F_st_ for molecular markers, Q_st_ is a metric measuring the population differentiation for phenotypic traits (Prout & Barker 1993; Spitze 1993). In theory, the Q_st_ of neutral traits on average should be equal to the mean F_st_ of neutral molecular markers (Rogers & Harpending 1992; Whitlock & Mccauley 1999; Whitlock 2008). We calculated P_st_ (Leinonen *et al*. 2006), a surrogate of Q_st_, based on the recombination rates of both linkage groups 8 and 9 (see Discussion for the implications of using P_st_). The mean P_st_ for recombination rate is 0.52 based on our ANOVA analyses of 1000 bootstrap replicates. We also estimated that the average genome-wide F_st_ of four-fold degenerate sites between *D. pulex* and *D. pulicaria* is 0.15 (0.12 and 0.20 for the interval on linkage group 8 and 9, respectively). Based on the distribution of F_st_ of genome-wide four-fold degenerate sites, we simulated the Q_st_ of a neutrally evolving trait to estimate the distribution of the test statistic Q_st_ – F_st_. Interestingly, our P_st_ – F_st_ value is significantly higher than the Q_st_ – F_st_ values of the simulated neutrally evolving trait (Fig. 3, P=0.03), leading us to reject the neutral hypothesis and to conclude that recombination rate divergence between these two species is adaptive.

**Figure 3.**
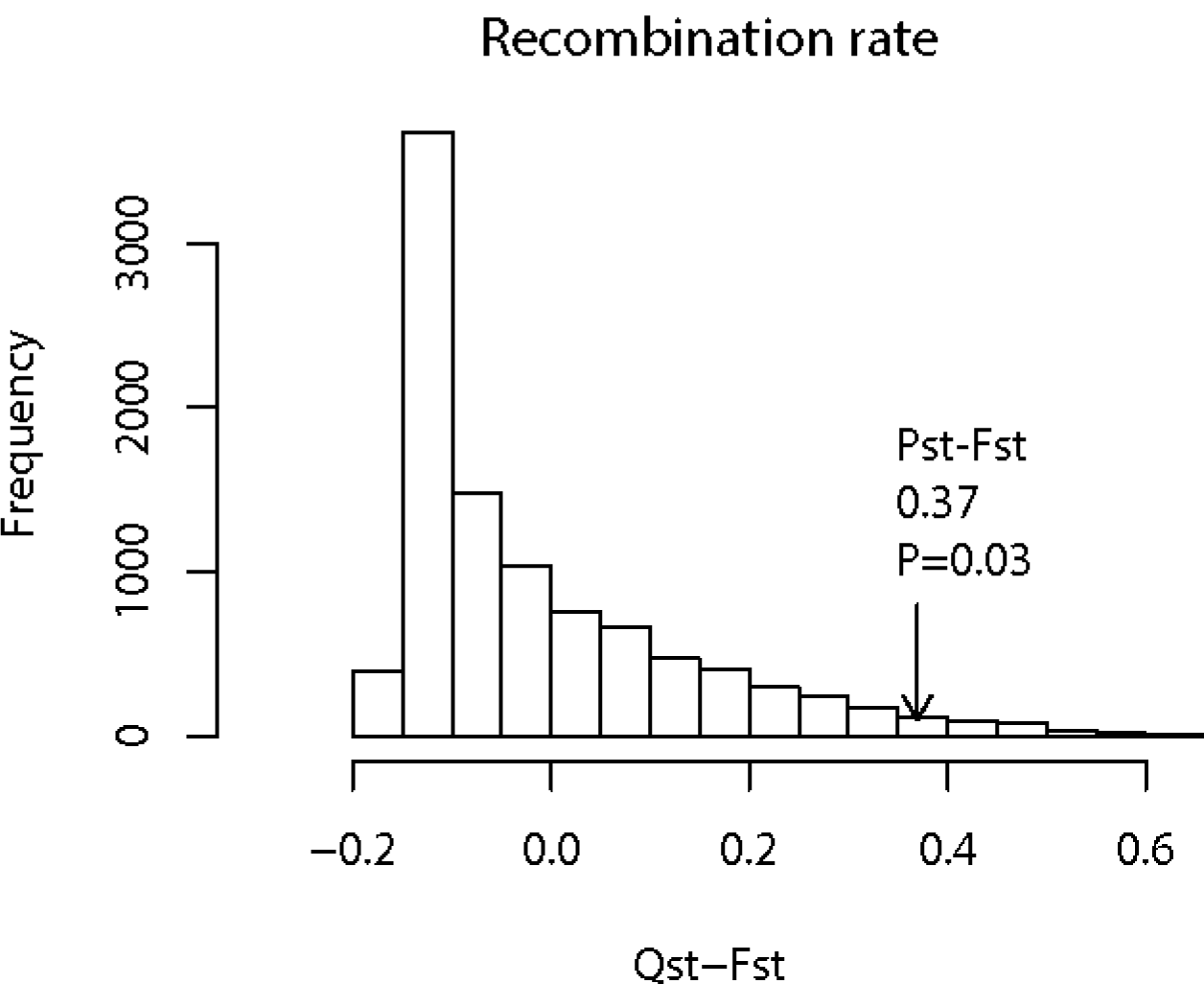
The simulated distribution of Q_st_ – F_st_ for a neutral trait and the observed P_st_ – F_st_ (indicated by an arrow).

## Discussion

### A new model system for studying recombination rate divergence

Meiotic recombination is one of the most laborious genetic parameters to estimate, with most species having no more than a handful of genetic maps each. Due to the lack of data on population-level recombination rate variation, many theories on the evolution of recombination remain untested in natural populations. In the past two decades only a handful of studies surveyed the recombination rate variation within and between populations from different environments. Prior work on the fungus *Sordaria fimicola* revealed heritable genetic variation in recombination rate between strains inhabiting harsh and mild habitat, with higher recombination rates found in the harsh habitat than in the mild environment (Saleem *et al*. 2001). Using novel single-sperm genotyping approach, the current study supports the hypothesis that directional selection coupled with habitat shift leads to elevated recombination rate in the model system of microcrustacean *Daphnia*. One could argue that the permanent lake habitat seems more stable and less harsh than ephemeral pond habitats and lower recombination rate would evolve in the lake species *D. pulicaria*. We suggest that stability of lake environment may not necessarily mean a benign environment for *Daphnia*. Other factors such as predator abundance may also determine the harshness of environment to *Daphnia*. Moreover, the higher recombination rate in *D. pulicaria* may result from the fact that *D. pulicaria* has lower frequency of sexual reproduction than *D. pulex* so that higher recombination rate evolves to produce genetically diverse offspring. Interestingly, based on P_st_ – F_st_ comparison analysis we find strong evidence that the divergence of recombination rates between these two species is adaptive and unlikely to be explained by genetic drift. With overall significantly higher recombination rates observed in *D. pulicaria* than in *D. pulex*, we argue that the directional selection that led to the local adaptation of *D. pulicaria* to permanent lake habitats (e.g., physiology, life history, see Introduction) most likely shaped the recombination rate divergence, providing support to the theory that directional selection leads to elevated recombination rate. However, with current data we cannot exclude the possibly that selection reduces recombination rate in *D. pulex*. We also suggest that the different recombination rates of the surveyed regions in these two species are unlikely due to translocation events (e.g., in one species the surveyed region moved from the tip of chromosome to a more central location). This is because the inter-specific F_1_s, F_2_s, and backcrosses have normal fertility (Heier & Dudycha 2009), suggesting no major chromosomal structure alterations affecting recombination.

It should be noted that different recombination rate between these two species is unlikely to be due to phenotypic plasticity as the habitat transition event most likely occurred ∼1-2 million years ago, and the examined *Daphnia* isolates have been acclimatized to lab conditions and were measured for recombination rates in a common garden experiment. On the other hand, as the current study is based on a relatively small set of populations and on two linkage groups, it remains to be seen whether the observed pattern holds true for the genome-wide recombination rate variation for a larger set of *D. pulex* and *D. pulicaria* populations.

Although prior work identified some empirical support for the theory that directional selection leads to elevated recombination rate, these work has largely been restricted to comparing domesticated animals, plants, and fungi (Ross-Ibarra 2004; Munoz-Fuentes *et al*. 2015; Saleem *et al*. 2001) with their wild progenitors and to examining laboratory populations (Korol & Iliadi 1994; Aggarwal *et al*. 2015). Although studies investigating intra- and inter-specific recombination rate divergence are not uncommon (reviewed in Smukowski and Noor (2011)), this kind of studies are usually deficient in an ecological understanding of the speciation process or lack the population-level sampling required for inferring the driving forces of recombination rate differentiation. However, a recent study examining two ecologically different populations of *Drosophila pseudoobscura* found that the divergence of genome-wide recombination rate is due to natural selection (Samuk *et al*. 2020). Our study is valuable in providing solid evidence in support of this hypothesis from the perspective of incipient species pairs undergoing ecological speciation. Notably, as *Daphnia* is different from the other examined species in that their sexual reproduction is triggered by environmental conditions (e.g., they only reproduce under certain environmental conditions and the frequency of sex is less frequent in *D. pulicaria* than in *D. pulex*), it is likely that recombination rate in *Daphnia* is subject to stronger selection to produce genetically diverse offspring than other species that engages in regular sexual reproduction. The lower frequency of sexual reproduction in *D. pulicaria* may also contribute to its higher recombination rate than *D. pulex*. We therefore argue that the well-understood ecology (distinct ephemeral pond vs. permanent habitats) and evolutionary history (speciation associated with transition from pond to lake habitats) of *D. pulex* and *D. pulicaria* set up an excellent framework for future in-depth investigation of the evolutionary and genetic basis of divergence in recombination rates.

### Single-sperm sequencing for studying recombination rate variation in emerging systems

A major hurdle in studying recombination rate variation is the laborious process of generating genetic maps. This study greatly benefited from the novel whole-genome sequencing technique developed for single-sperm cells (Xu *et al*. 2015; Xu & Young 2017). Single sperm sequencing emerged in 1980’s as a methodology for estimating localized recombination rates (Li *et al*. 1988; Cui *et al*. 1989). Nonetheless, empowered by whole-genome sequencing technologies, this technique has recently been applied to human and mouse to examine whole-genome recombination patterns (Wang *et al*. 2012; Lu *et al*. 2012; Hinch *et al*. 2019).

We note that as collecting a large number of sperm/pollen cells is feasible in many species, our experimental procedure for single-sperm sequencing/genotyping can be applied to other emerging model systems. Even for species with large genome sizes, we have seen multiple studies using single sperm sequencing to examine the regulation of recombination events in mouse (Hinch *et al*. 2019) and human (Bell *et al*. 2020). Although our protocol uses flow cytometry to isolate single cells and relies on whole-genome amplification of single cells, these are currently common laboratory procedures. We hope that an increasing number of researchers will take advantage of this approach to examine the divergence of recombination rates in a diverse set of emerging model systems with interesting ecological attributes. Nonetheless, the sperm sequencing approach does not allow to evaluate sexual dimorphism and fine-scale variation in recombination rate.

### P_st_-F_st_ comparison

We used P_st_-F_st_ comparison to determine whether the divergence of recombination rate between *D. pulex* and *D. pulicaria* is adaptive. This test is based on the observation that for neutral phenotypic traits that are controlled by purely additive genes the mean Q_st_ (we used P_st_ as a surrogate for Q_st_) is equal to the mean F_st_ of neutral genetic loci (Lande 1992; Whitlock 1999). While *Q*_*st*_ for a quantitative trait is calculated using additive genetic variance obtained through breeding experiments, *P*_*st*_ is based on total phenotypic variance, which could be inflated due to environmental factors, thus complicating the interpretation of P_*st*_-F_*st*_ comparison. Although the observation Q_st_=F_st_ for a neutral trait is based on several assumptions and this study likely violated some of them, we argue that the strong evidence pointing to the adaptive nature of the observed divergence in recombination rate is unlikely compromised (see below).

An important assumption of Q_st_=F_st_ for neutral phenotypic traits is that the loci from which F_st_ is derived should be neutral. Although there have been concerns about whether Q_st_=F_st_ when the F_st_ is based on markers such as microsatellites that have high mutation rate (Hendry 2002), the use of SNPs in our study alleviates this concern. Furthermore, despite that in other species four-fold degenerate sites have been shown to experience purifying selection, population genomic analyses of *D. pulex* show that these sites evolve in a nearly neutral fashion (Lynch *et al*. 2017). Therefore, our use of genome-wide four-fold degenerate sites (n= 94711) should provide a meaningful estimate of the mean F_st_ of neutral sites.

Our analysis differs from the standard Q_st_-F_st_ analysis in the use of P_st_ as a substitute of Q_st_. However, this study differs significantly from studies directly collecting phenotypic data from the field because the recombination rates were estimated in a common garden experiment. As recombination rate is known to be of great phenotypic plasticity due to biotic and abiotic factors such as age and temperature (Lloyd *et al*. 2018; Hunter, Robinson, *et al*. 2016), we estimated recombination rates from 2-week old males that were maintained under controlled temperature and photoperiod. Therefore, the obtained P_st_ value is unlikely to be inflated by environmental effects.

Because Q_st_ is defined based on additive variance of traits, one may wonder whether the P_st_ – F_st_ test in this study is biased towards rejecting the neutral hypothesis. Based on previous work that examines how dominance and epistatic effects may affect this test, we argue that our results are unlikely to be biased. Although the genetic basis of recombination rate variation (e.g., relative contribution of additive variance, dominance effects, and epistasis) is poorly understood, we consider the potential impact of epistasis and dominance effects in turns. It is true that our P_st_ estimates could be affected by dominance and epistasis. However, it has been shown that epistasis tends to produce Q_st_ values less than neutral F_st_ (Whitlock 1999). Similarly, dominance makes Q_st_ equal or less than neutral F_st_ under the assumption of an island model (Goudet & Büchi 2006; Goudet & Martin 2007). Even though dominance under limited demographic circumstances can make Q_st_ of neutral traits exceed neutral F_st_, this is unlikely for traits affected by multiple loci (Goudet & Martin 2007) such as recombination rates. Taking all these into consideration, we argue that our use of P_st_ in this study makes our test likely conservative.

Due to limited resources this study only examined the recombination variation of two genomic regions in males. Despite the promising evidence that the positive selection is responsible for the divergence in recombination rate between *D. pulex* and *D. pulicaria*, it remains unclear whether this is true for the genome-wide variation in males and whether this is true for female-specific recombination rate. Sex-specific difference in recombination rates often shows that females recombine more frequently than males (see Sardell & Kirkpatrick 2020). For example, human females on average recombine 1.6 times as much as males (Kong *et al*. 2010), and sticklebacks show a similar pattern (Sardell *et al*. 2018). It will be interesting to examine the female recombination rate divergence between *D. pulex* and *D. pulicaria* and determine whether male and female recombination rate evolution is shaped by the same evolutionary forces.

### Within-species recombination rate divergence

With much of the focus of this study probing whether between-species divergence in recombination rate is adaptive, it is necessary to provide some explanation about the contrasting pattern of within-species divergence in these two species. As mentioned in the Results, the intra-specific recombination rate of *D. pulex* varies by nearly 3 fold for both linkage groups 8 and 9, whereas the intraspecific variation within *D. pulicaria* is much lower with a ∼1.3 fold difference on linkage group 8 and little variation on linkage group 9 (Figs. 1 and 2). The within-species divergence in *D. pulex* is larger than all the currently available within-species divergence (reviewed in Ritz *et al*. 2017), such as ∼1.6-fold variation in both sexes of human (Coop *et al*. 2008), 1.9 fold in mice (Dumont *et al*. 2009), 1.1-2 fold in *Drosophila* (Brooks & Marks 1986; Hunter, Huang, *et al*. 2016), 1.3 fold in *Arabidopsis* (Sanchez-Moran *et al*. 2002), whereas the within-species divergence in *D. pulicaria* is in line with these available estimates.

One plausible explanation for this drastic difference between these two species is that selection pressure for maintaining recombination rates among different *D. pulicaria* populations is much more uniform than among *D. pulex* populations. To better understand this, we can use results of previous work on how spatially heterogeneous selection pressure influences the evolution recombination rate (Lenormand & Otto 2000). Regardless of the forms of epistasis and linkage disequilibrium and the amount of linkage between recombination rate modifier and the selected loci, when environmental selection pressures vary between populations with frequent migration it is predicted that more variation in recombination rate is expected in populations inhabiting highly spatially variable environments (Lenormand & Otto 2000). Although it is often said that the typical habitat of *D. pulex* is ephemeral pond habitats, we have to acknowledge that ecological conditions of each pond population probably differ substantially in terms of pond sizes, depths, hydrological conditions, habitat heterogeneity, predators, and other biotic and abiotic factors. On the other hand, the ecology of the different stratified permanent lake habitats of *D. pulicaria* may differ to a lesser extent. We therefore hypothesize that the greater variability of recombination within *D. pulex* is likely due to the greater amount of heterogeneity among the pond habitats. This hypothesis is certainly worth future investigation by examining a large number of populations of each species, which can provide insight into how spatially heterogeneous selection shapes the evolution of recombination rates.

## Materials and Methods

### Daphnia culture and sperm extraction

Males were collected for three isolates of *D. pulex* and *D. pulicaria* each (Table 1). Each isolate represents a distinct population and clonally produced males (i.e., genetically identical excluding rare mutations) of each isolate were collected. To avoid maternal effect on recombination rate, females of each isolates were maintained in the same conditions for 2 generations. To induce the clonal production of males, mature females of the third generation with early sign of carrying broods were collected and cultured at 20 °C in artificial lake water (Kilham *et al*. 1998) containing 400 nM Methyl Farnesoate (MF), a juvenile hormone that determines the sex of *Daphnia* offspring (Olmstead & Leblanc 2002). They were fed *ad libitum* with a suspension of *Scenedesmus obliquus*, and the offspring were screened for males. A total of 15-18 males were collected from each clone (same genotype) and were maintained in the lab for two weeks before sperm collection.

**Table 1:** Summary of the *Daphnia* isolates used for the recombination rate estimates, with sampling locations and NCBI SRA accession numbers for whole-genome sequencing raw reads.

| Species | Isolates | SRA (NCBI) | Lab code | Location |
| --- | --- | --- | --- | --- |
| *D. pulex* | px1 | SRX4386564 | SW4 | Illinois |
|  | px2 | SRX4386576 | LPB17 | Long point, Ontario, Canada |
|  | px3 | SRX4386574 | Tex21 | 42°12, -83°12, Textile Road, Michigan |
| *D. pulicaria* | pa1 | SRS1024794 | Little Curtis | 45°43, -122°44, Curtis Lake, Oregon |
|  | pa2 | SRS1024791 | RLSD26 | 44°57, -96°49, Round Lake, South Dakota |
|  | pa3 | SRS1024797 | AroMoose | 44°50, -69°16, Sebasticook Lake, Maine |

For analyzing recombination rate of each *Daphnia* isolate, we collected sperm from all the identified males because they have identical genotype. To extract sperm, each male immersed in a drop of double-distilled water (ddH_2_O) was gently pressed with a cover-slip on a microscope slide. The ddH_2_O surrounding each individual was collected using Sigmacote-washed capillary needles and mouth pipettes into a 1.5 mL microcentrifuge tubes containing 50 μl of 1x PBS solution (Xu *et al*. 2015). To facilitate the sorting of single sperm cells by flow cytometry, we stained sperm cells using 8 μl of Hoechst 33342 (100 μg/mL) (Sigma-Aldrich) and incubated the sample in the dark at room temperature for 2 hours.

### Single-sperm cell sorting

A BD FACS Aria-II cell sorter was used to isolate single sperm cells into individual wells of 96-well PCR plates containing cell lysis buffer. The specific settings of the FACS Aria II instrument were 488 nm 100 mW laser for light scatter detection and 355 nm 20 mW for Hoechst detection. A nozzle of 70 mm was used at 45 psi, and FSC-PMT was used for optimal small particle discrimination.

Each well of the PCR plate contained 5 μl of lysis buffer consisting of Tris (30 mM), EDTA (2 μM), potassium chloride (20 μM), Triton (0.2%), DTT (40 mM), and protease/ Proteinase K (2.5 μg/μl). Cell lysis was performed in a thermal cycler at 50 °C for 3 hours, 75 °C for 20 minutes, and 80 °C for 5 minutes.

### Whole-Genome Amplification

To obtain enough DNA from each sperm for genotyping, the lysed single sperm cell was used for MALBAC (Multiple annealing and looping-based amplification) whole-genome amplification (Zong *et al*. 2012). MALBAC consists of a pre-amplification stage and a standard PCR amplification. The pre-amplification is initiated with random primers, each having a common 27-nucleotide sequence (5□-GTGAGTGATGGTTGAGGTAGTGTGGAG-3□) and 8 variable nucleotides that can evenly hybridize to the templates.

### Pre-amplification stage

A solution of 3.0 μl ThermoPol buffer (New England Biolabs), 1 μl dNTPs (10 mM), 0.75 ul each of two primers NT and NG (10 μM), and 19.5 μl H_2_O was added to each sperm sample. The samples were incubated at 95 °C for 5 min and quenched immediately on ice. 0.5 μl of *Bst* large fragment polymerase (New England Biolabs) was added to each sample and the following thermal amplification regime is performed: 10 °C for 45 sec, 15 °C for 45 sec, 20 °C for 45 sec, 30 °C for 45 sec, 40 °C for 45 sec, 50 °C for 45 sec, 65 °C for 2 min, 95 °C for 20 sec, followed by quenching on ice. Subsequently, five cycles of pre-amplification cycles were performed, consisting of 10 °C for 45 sec, 15 °C for 45 sec, 20 °C for 45 sec, 30 °C for 45 sec, 40 °C for 45 sec, 50 °C for 45 sec, 65 °C for 2 min, 95 °C for 20 sec, and 58 °C for 40 sec, followed by quenching on ice. 0.5 μl *Bst* large fragment polymerase was added to each sample before carrying out the next cycle.

### Standard PCR amplification stage

A standard PCR amplification was performed on the amplicons from the pre-amplification stage using the 27mer as primer (5□-GTGAGTGATGGTTGAGGTAGTGTGGAG-3□) to generate the 1-2 μg DNA required for downstream genotyping. Each reaction consisted of the product from the pre-amplification, 3 μl ThermoPol Buffer (New England Biolabs), 1 μl dNTPs (10 mM), 23.5 ml H_2_O, 1.5 μl 27mer (10 μM), and 1 μl DeepVentR exo-polymerase (New England Biolabs). The PCR thermal regime consisted of 22 rounds of 94 °C for 20 sec, 59 °C for 20 sec, 65 °C for 1 min, 72 °C for 2 min, which was followed by 72 °C for 5 min.

### Recombination rate estimation

To examine recombination rate variation in *D. pulex* and *D. pulicaria*, we focused on two regions that are at the tip of the linkage groups and have ∼20 cM genetic distance on linkage groups 8 and 9 from the microsatellite-based genetic map by Cristescu *et al* (2006). For linkage group 8, located between the microsatellite markers d077 and d068, the interval is 1.5 Mbp, whereas on linkage group 9, the region spans ∼0.5 Mbp lying between the microsatellite markers d171 and d118.

For detecting recombination events, two heterozygous markers are required. However, the four mapped microsatellite markers (i.e., d077, d068, d171, d118) are not heterozygous in all the *Daphnia* isolates. In cases where any of these markers are homozygous in any isolate, new heterozygous microsatellites were identified within a 50-kb window centered at the mapped marker and were used for estimating recombination (Supplementary Table 1). The web-based platform WebSat (Martins *et al*. 2009) was used for identifying microsatellite markers and primer designs.

Our microsatellite genotyping followed the strategy outlined by (Schuelke 2000). Briefly, a M13 tail is added to the 5’ prime end of the forward primer, and a M13 sequenced labelled with one of the NED, PET, FAM and VIC fluorescent dye was used in the PCR. The thermal cycling program for microsatellite amplification consisted of 3 min at 95 °C, 10 cycles of 35 sec at 95 °C, 35 sec at 56 °C (the temperature increased by 1 °C for each cycle) and 45 sec at 72 °C, 30 cycles of 35 sec at 95 °C, 35 sec at 48 °C, 45 sec at 72 °C, and a final 10 min at 72 °C. Fragment analysis was performed on an ABI 3130 Genetic Analyzer (Life Technologies) using 20x diluted PCR product. The four different M13 dyes allowed the pooled genotyping of different markers labelled with different dyes. The genotypes were called using the software GeneMapper 4.0 (Life Technologies).

To estimate the recombination rate for the two intervals of interests, 2-locus genotypes were examined for the pool of genotyped sperm for each *Daphnia* isolate. The number of sperm genotyped for each *Daphnia* isolate ranged from 73-94. The two most abundant genotypes were identified as the parental genotypes, whereas the two rare genotypes were derived from recombination events. For example, the two locus genotypes for d077 (alleles: 227 and 232 bp) and d068 (alleles: 337 and 343 bp) are 227/337 (10 sperm cells), 232/343 (10 sperm cells), 232/337 (40 sperm cells), and 227/343 (40 sperm cells). Then, the genotypes 227/337 and 232/343 are recognized as recombinant genotypes with a recombinant frequency of 0.2. The frequency of recombinants is converted to Kosambi centiMorgan map distance. The standard error (SE) of recombination was calculated as 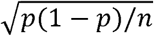, where the p represents proportion of recombinant sperm cells and n represents the number of sampled sperm cells.

### P_st_-F_st_ comparison

To investigate whether the divergence of recombination rate between these two species is adaptive, we performed P_st_-F_st_ comparison analysis. As the divergence of quantitative traits can be shaped by mutation, selection, and genetic drift, various methods have been developed for deciphering whether the divergence of phenotypic traits is neutral (i.e., can be adequately explained by drift alone) or adaptive. An important approach among these is the comparison of Q_st_ and F_st_ values.

Analogous to the famous F_st_ for measuring population differentiation using molecular markers (reviewed in Holsinger and Weir, (2009)), Q_st_ (Prout & Barker 1993; Spitze 1993) is established as a measure of the genetic differentiation among populations for phenotypic traits. For a neutral quantitative trait with additive genetic basis, its Q_st_ value on average should be equivalent to the mean F_st_ of neutral loci (Rogers & Harpending 1992; Whitlock & Mccauley 1999; Whitlock 2008), providing an important means for distinguishing between neutral and adaptive divergence. Therefore, if the Q_st_ of a trait is significantly higher than the mean F_st_ of neutral loci, it would indicate divergent selection on this trait. On the contrary, if Q_st_ of a trait is significantly smaller than the mean F_st_ of neutral loci, it would indicate stabilizing selection on the trait in the presence of drift. Moreover, identical values of Q_st_ and F_st_ would indicate no evidence for selection acting in a spatially heterogeneous manner.

As specific breeding experimental designs in a common garden environment are required for estimating additive variance that is required for calculating Q_st_, many studies on wild populations used another metric P_st_ that is a surrogate to Q_st_ (Leinonen *et al*. 2006). P_st_ is a metric measuring total phenotypic variance (rather than additive variance) among populations, which could be confounded by environmental effects for phenotype data directly collected from the field. Although our recombination rates were measured in a controlled environment and in same-aged males, our experiments did not allow us to estimate the additive variance. Thus, we decided to use P_st_ as a surrogate for Q_st_ in this analysis.

To estimate P_st_ of recombination rate, we used recombination rate data for both chromosomes 8 and 9 to quantify within- and between-species variances using ANOVA in R. This strategy gave us a larger sample size and more statistical power than examining single linkage groups alone. The between-species variance was calculated using the equation Var(s) =(MSs – MSe)/n, where MSs and MSe represent the mean squares of between- and within-species, respectively, and n represents the number of data points for each species (n=6). The within-species variance Var(e) is equal to MSe, which is the mean squares of within-species. The P_st_ value is calculated using the equation Var(s)/[Var(s) + 2Var(e)]. A total of 1000 bootstrap replicates were generated and analyzed using ANOVA to estimate the distribution and mean value of P_st_.

To determine whether the divergence of recombination rates between *D. pulex* and *D. pulicaria* is adaptive, we followed the approach of (Whitlock & Guillaume 2009) to examine the difference between Qst and Fst with Qst – Fst as the test statistic. This approach rests upon the notion that the mean Q_st_ value of neutral traits is expected to be the same as the mean F_st_ of neutral makers under certain assumptions (Whitlock & Guillaume 2009). The Fst between *D. pulex* and *D. pulicaria* was estimated using genome-wide four-fold degenerate sites (n=94711) extracted from the whole-genome sequences of these isolates from (Tucker *et al*. 2013).

To simulate the distribution of the Q_st_ of a neutral trait, we calculated the expected between-species variance Var(s) using the formula Var(s) = 2F_st.bootstrap_·Var(e)/(1-F_st.bootstrap_), where F_st.bootstrap_ is the mean value of a bootstrap sample of four-fold degenerate sites and Var(e) is the observed within-species variance. Then we simulated the between- and within-species variance, Var(s).hat and Var(e).hat, respectively. Var(e).hat was calculated as ^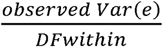^ multiplied by a random number drawn from a chi-square distribution with the degree of freedom at within-species level (i.e, DFwithin), whereas Var(s).hat was simulated as ^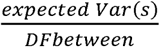^ multiplied by a random number drawn from a chi-square distribution with the degree of freedom at between-species level (i.e., DFbetween). Furthermore, the simulated Q_st_ was calculated as Var(s).hat/[Vars(s).hat + 2Var(e).hat]. The simulation was repeated for 10,000 times to obtain a distribution of the test metric Q_st_ – F_st_. Lastly, we determined whether the observed P_st_ – F_st_ differs significantly from the neutral expectations by identifying the quantile of simulated distribution that had higher values than the observation, which gave us the P value of the test. This procedure was perform using a R script slightly modified from Lind *et al*. (2011).

## Acknowledgements

We thank Marelize Snyman, Trung Huynh, Hongjun Wang, and Thinh Pham for their help and constructive comments on this manuscript. This work is partly supported by start-up funds from University of Texas at Arlington and is partly supported by NIH grant R35GM133730 to SX.

