## Supplementary Table 1 for "Adaptive divergence of meiotic recombination rate in ecological speciation"

**Supplementary Table1**. Summary of microsatellite markers genotyped in each isolate for linkage groups 8 and 9.

| **Species** | **Isolate** | **Linkage Group** | **Markers** | **Forward Primer(5ʹ-3ʹ)** | **Reverse Primer(5ʹ-3ʹ)** |
| --- | --- | --- | --- | --- | --- |
| *D. pulex* | px1(SW4) | 8 | d077 | TGTGGAACAGATGGCGACTA | CACTCTCAACGATCCAAGCA |
|  |  | 8 | d068-2 | ATATTTCCACCGACGTTTTCAC | TGTTGTATCCAGTTGCCTTTTG |
|  |  | 9 | d171-1 | TGTCCGTCTCTACTTCCATTCA | CCCTTATTTTCTCCCGTGTGTA |
|  |  | 9 | d118 | ACTCGACACAAGCGGAAAGT | AAAGGGAGGAGCTGAAATCC |
| *D. pulex* | px2(LPB17) | 8 | d077-6 | TAAAATACACACACACGCAGCA | CAGTAGTTCCCTCAACTCGCTT |
|  |  | 8 | d068-2 | ATATTTCCACCGACGTTTTCAC | TGTTGTATCCAGTTGCCTTTTG |
|  |  | 9 | d171-3 | CGGTCCTCGGTAGACATTTAGT | GTTGTCATTACCCCGATCCTT |
|  |  | 9 | d118-3 | ATCTGCTTTCATGGTGCTCTTT | TAGCGCGGGTTTCATAATAACT |
| *D. pulex* | px3(Tex21) | 8 | d077-5 | ATGAGAATACGGCCACCTTG | CGTTTACGACCTCGCATAAAGT |
|  |  | 8 | d068-2 | ATATTTCCACCGACGTTTTCAC | TGTTGTATCCAGTTGCCTTTTG |
|  |  | 9 | d171-3 | CGGTCCTCGGTAGACATTTAGT | GTTGTCATTACCCCGATCCTT |
|  |  | 9 | d118-8 | AGTTATTATGAAACCCGCGCTA | GGTGAGAAATTGTGTTCGTTCA |
| *D. pulicaria* | pa1(LittleCurtis) | 8 | d077-4 | AATCCTTATTGCACAGCCTCAT | CGTTCCTTTTCTTCATTTCCAG |
|  |  | 8 | d068-2 | ATATTTCCACCGACGTTTTCAC | TGTTGTATCCAGTTGCCTTTTG |
|  |  | 9 | d171 | AAGGACGACATCTGGCAATC | AATCGATCAGAACCGACACC |
|  |  | 9 | d118-1 | TAATAATGTGTGTAACCGCGCA | TTGATTTTCCTGGTGGTGGTAT |
| *D. pulicaria* | pa2(RLSD26) | 8 | d077-4 | AATCCTTATTGCACAGCCTCAT | CGTTCCTTTTCTTCATTTCCAG |
|  |  | 8 | d068-2 | ATATTTCCACCGACGTTTTCAC | TGTTGTATCCAGTTGCCTTTTG |
|  |  | 9 | d171-8 | CGTGAGTCCTTTCCGTGATATT | GCTTTCGATTTGTTCTTGCACT |
|  |  | 9 | d118-1 | TAATAATGTGTGTAACCGCGCA | TTGATTTTCCTGGTGGTGGTAT |
| *D. pulicaria* | pa3(AroMoose) | 8 | d077 | TGTGGAACAGATGGCGACTA | CACTCTCAACGATCCAAGCA |
|  |  | 8 | d068-2 | ATATTTCCACCGACGTTTTCAC | TGTTGTATCCAGTTGCCTTTTG |
|  |  | 9 | d171-8 | CGTGAGTCCTTTCCGTGATATT | GCTTTCGATTTGTTCTTGCACT |
|  |  | 9 | d118-7 | ACTTGAACCCATCGAGAAGGTA | CCTTGAGTTTGGACGTGACTTT |
